## supplementary data for "The satiety hormone cholecystokinin gates reproduction in fish by controlling gonadotropin secretion"

**Supplementry**

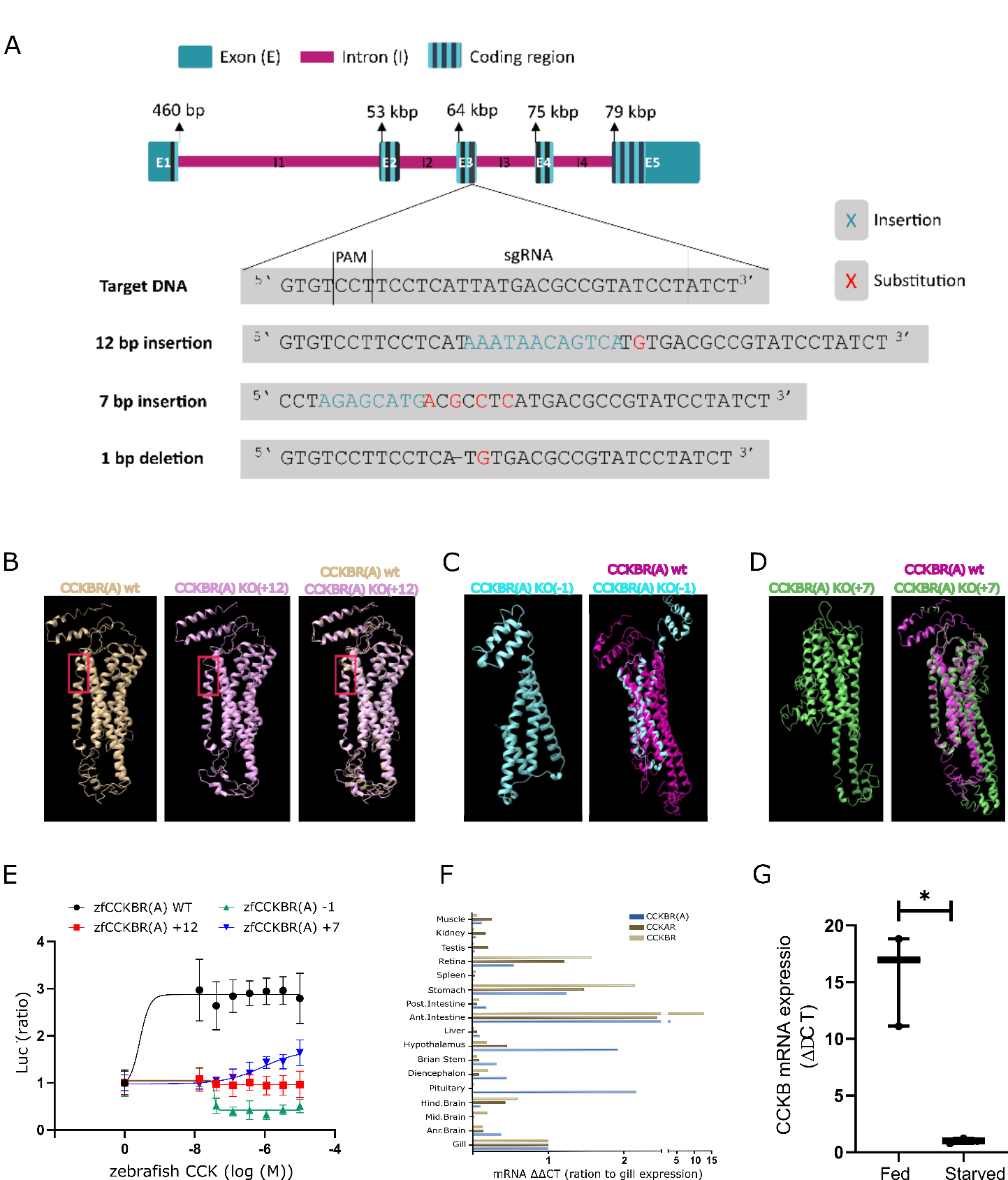

Figure 1-Figure supplement 1.

(A) The three identified mutations induced by gRNA number two. The 7 bp insertion and 1 bp depletion created a frameshift that disrupt the translation of the RNA, while the 12 bp insertion added four amino acids that disrupt the structure of the CCKR transmembrane domain of the receptor. (**B)** The protein structure of the CCKR with the +12 insertion (pink) compared to the WT CCKR structure (brown). The +12 addition resulted in the incorporation of four additional amino acids, specifically two asparagine, one serine, and one histidine, at residue number 187-191. Notably, this mutation-induced alteration is evident in the figure, where the WT CCKR displays an intact alpha helix structure and the mutant CCK receptor reveals a loss of this structural element. (**C)** The protein structure of the CCKR with a -1 nucleotide depletion (cyan) results in a frameshift, resulting in addition of a premature stop codon at residue 187, terminating the translation at this point, compared to the 442 amino acids in the WT CCKR (magenta). (**D)** The protein structure of the CCKR with the addition of 7 nucleotides at residue 174 (green), resulting in a frameshift and the introduction of a stop codon, thus yielding a receptor composed of 184 amino acids, contrasting the 442 amino acids in the WT receptor (magenta).All structures were created using the I-TASSER protein structure algorithm. **(E)** The CCK ligand stimulated WT CCKR but not the mutated receptors in a COS-7 reporter assay. Data are presented as mean ± SEM of a representative experiment performed in triplicate. **(F)** Expression levels by real-time PCR of CCKAR, CCKBR and CCKBRA in various tissues of Nile tilapia Fish. The relative expression of mRNA in each tissue was normalized to the expression levels of *ef1a* by the comparative threshold cycle method, all expression levels were then normalized to the expression levels of the gill for each gene respectively. (**G**) CCKB mRNA expression in the brain is significantly down-regulated in starved fish compared to fed fish (n=3, Paired t test, *p <0.05).

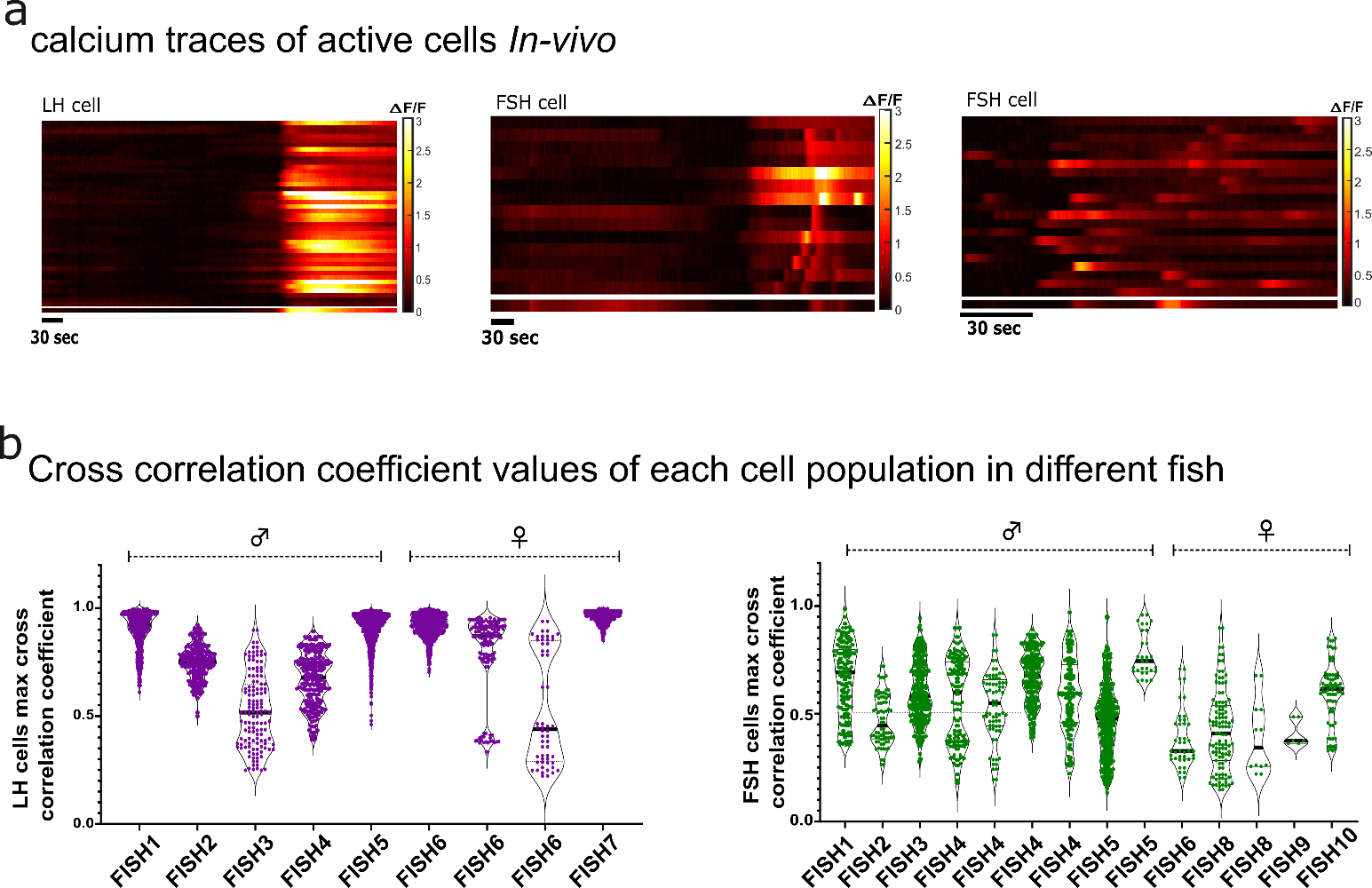

A

B

Figure 2- Figure supplement 1

**(A)** Heatmap of calcium traces from a pituitary imaged in vivo. Each line represents a cell. **(B)** Graphs showing repeated measures of max cross-correlation coefficient values of LH and FSH cells calcium activity.

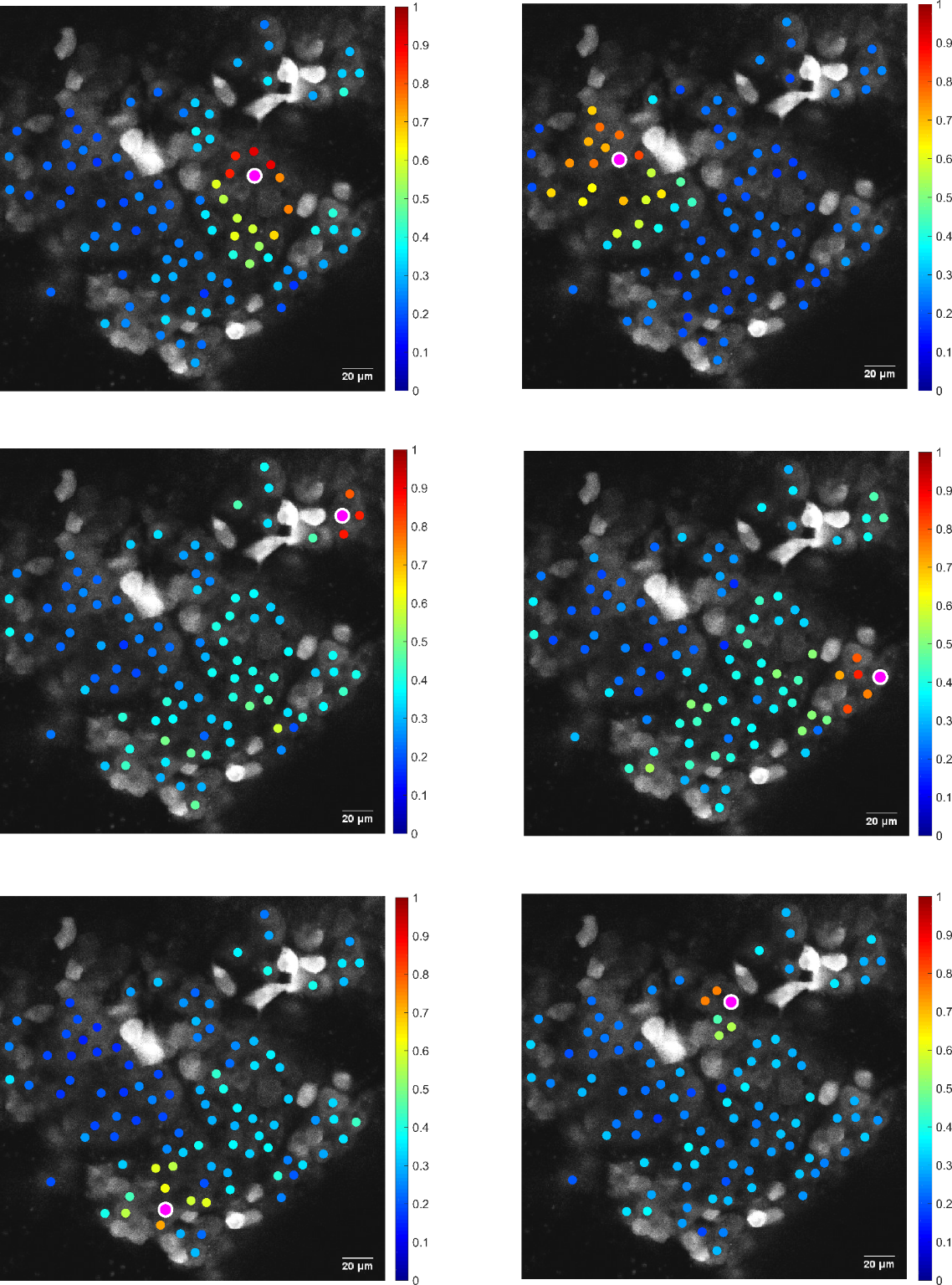

Figure 3 - Figure supplement 1

Calcium analysis of basal activity in LH cells reveals the synchronized spontaneous activity of small clamps of cells. Color-coded data points of cross-correlation coefficients between all cells and a chosen RIO (pink dot). In each panel, a different ROI was chosen, and a few nearby cells with high cross-correlation values are seen.

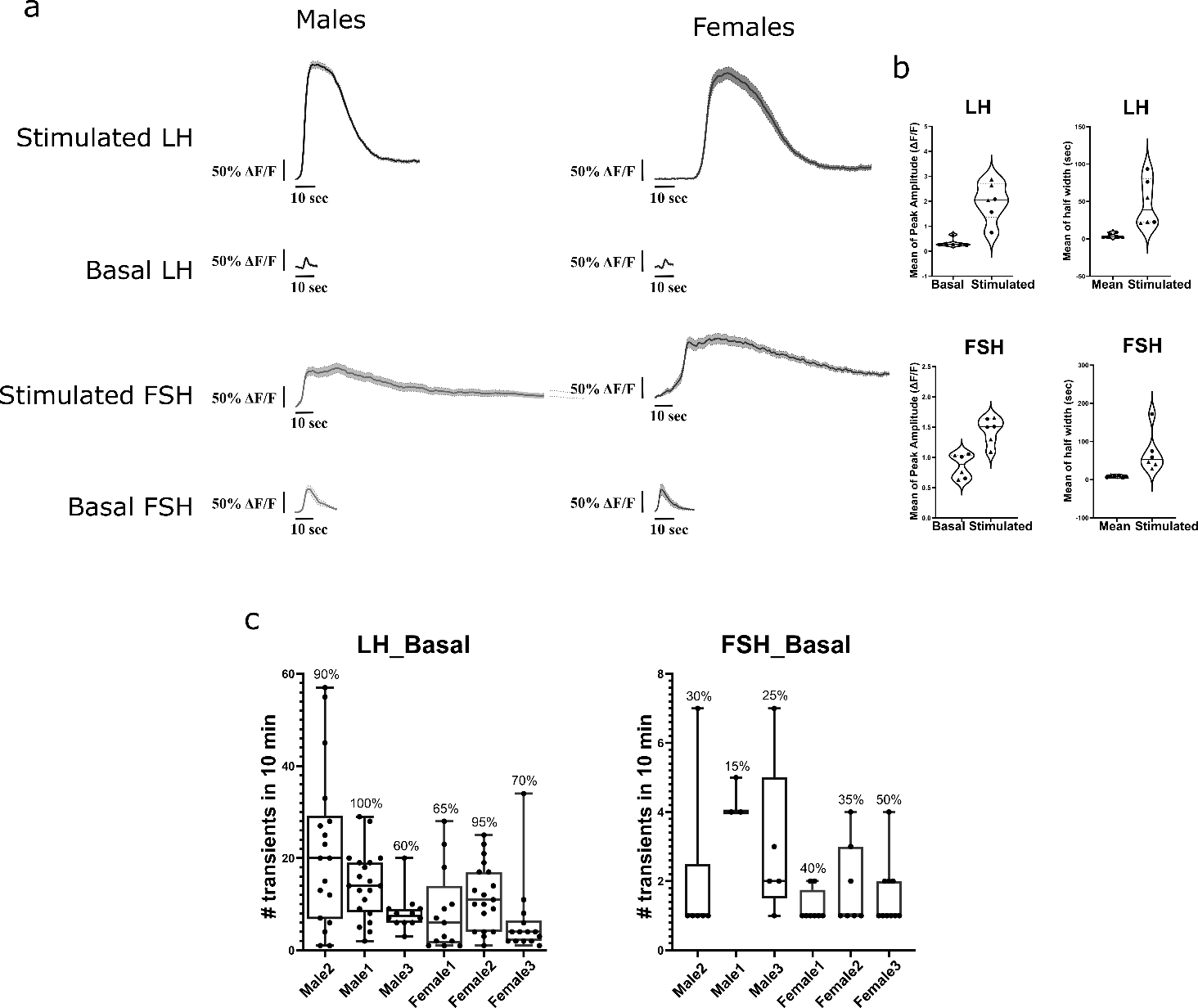

A

B

C

Figure 3 - Figure supplement 2.

**(A)** Examples of basal and stimulated calcium transients in LH and FSH cells of males and females. All transients are in the same scale of 50% ΔF/F and 10 sec. **(B)** The mean peak amplitude and half-width of each cell type, reflecting the intensity and duration of transients. Each dot represents the mean of 20 cell traces in a single female (circles, n=3) or male (triangles, n=3) fish. **(C)** The number of identified transients in basal traces during a 10-minute interval. On the top of each column is the percentage of active cells (i.e., cells in which transients were identified). Analysis was performed using pCLAMP 11 (molecular devices).

**Figure 4- Figure supplement 1.**

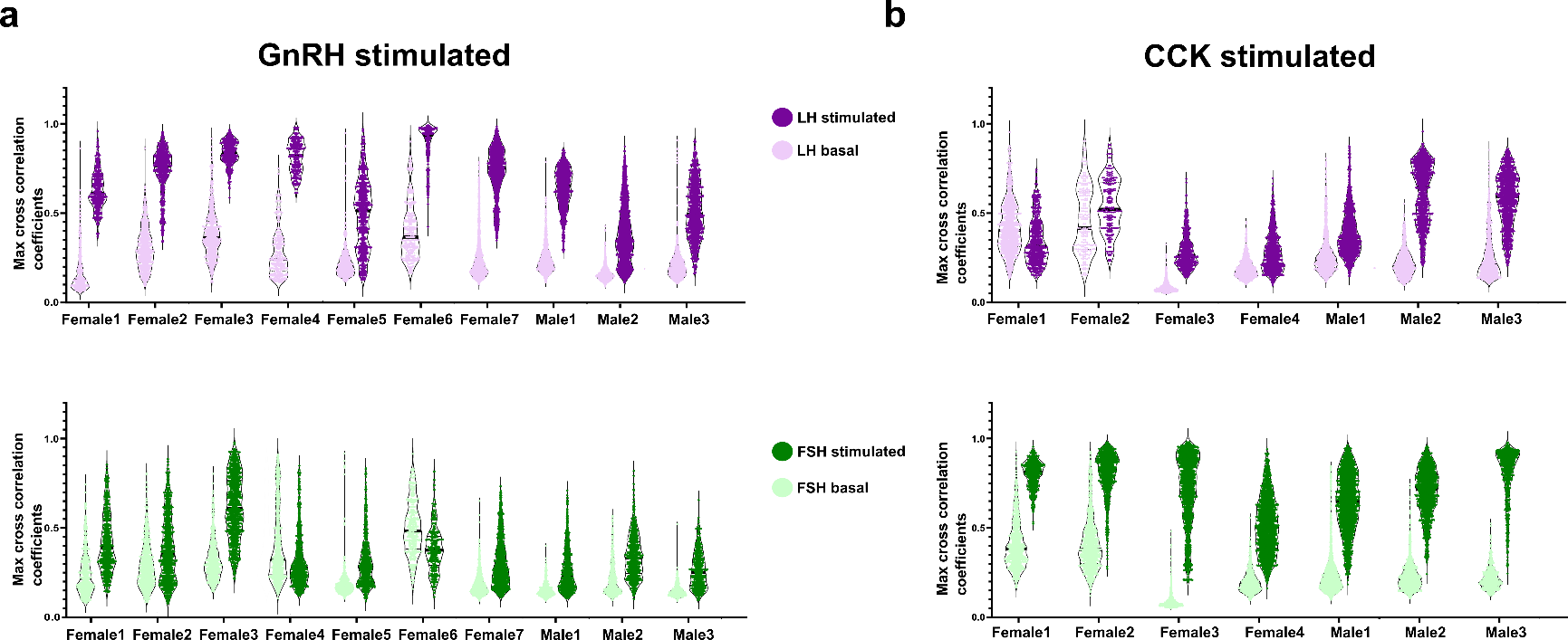

A

B

Max cross-correlation coefficient values between the calcium activity of all measured LH (magenta) or FSH (green) cells in each fish. **(A)** Compared basal activity to GnRH stimulated activity. **(B)** Compared basal activity to CCK stimulate activity.

Male

Female

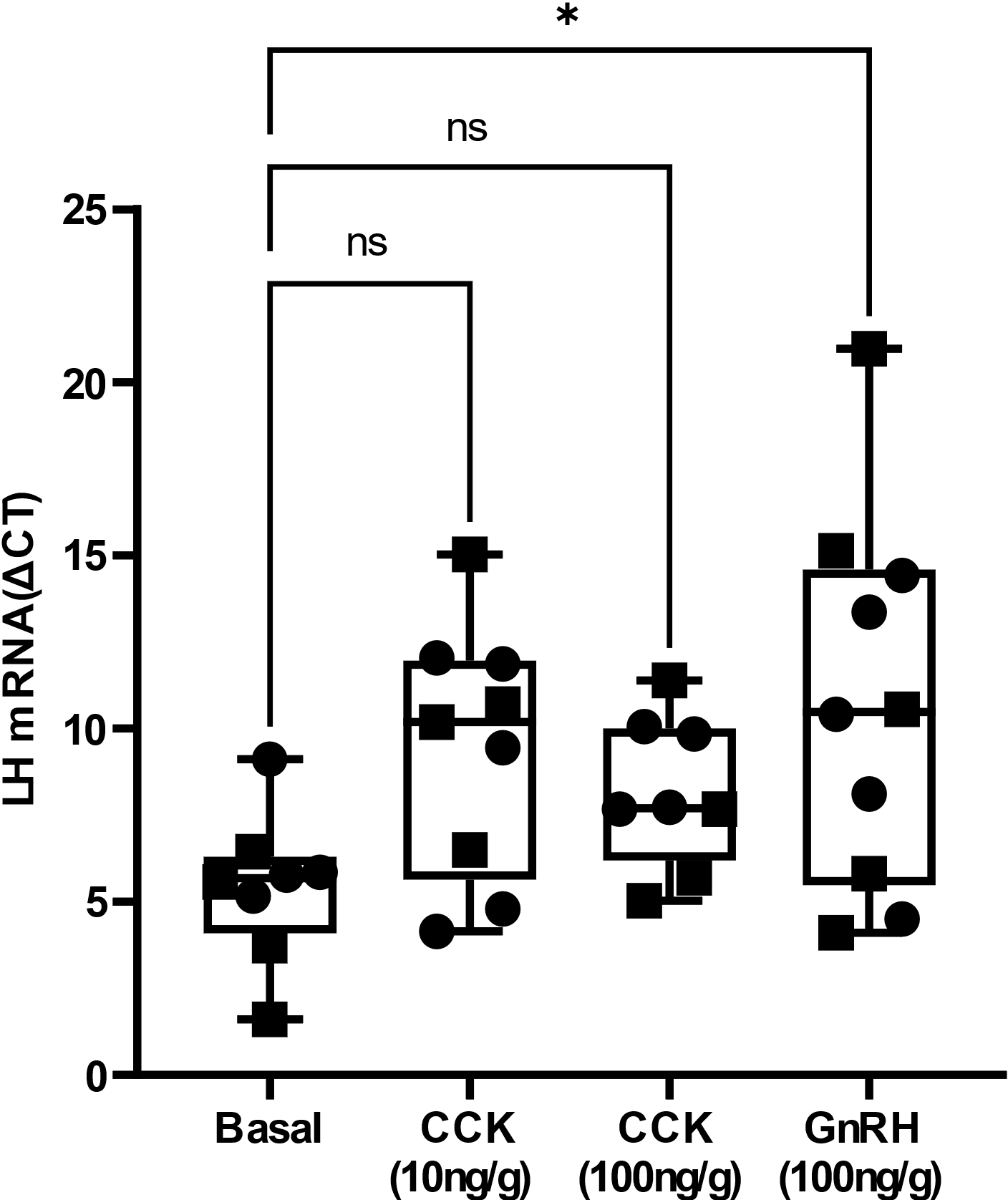

Figure 5- Figure supplement 1.

LH expression in the pituitary after in vivo injection of different concentration of CCK and GnRH. Expression level increased significantly only in response to GnRH injection.

| **Guide #** | **Sequence (5’ 🡪 3’)** | **Primers for HRM screening**  **Sequence (5’ 🡪 3’)** |
| --- | --- | --- |
| 1 | ACAACGCGTCGATTCTGAAC | Forward: AACAGTTGTGGTCAGGTCCG; Revers: CGCATTTGACAGAAAGCGCA |
| 2 | TAGGATACGGCGTCATAATG | Forward: TGCTTTGCGAGTGAAATGCTGA; Revers: GCCTTCTCCACCTCGCTATTCT |
| 3 | CCCAACAGGCTCCGCAAAGG | Forward: TCTTCTGCTACCTCACAAGCACT; Revers: CGGATCACTCGCTTCTTGGC |
| **Primers for real time PCR of tissue distributions, sequence (5’ 🡪 3’)** | | |
| TiCCKAR-F | CAAGGTCATCACTGCCACCT |  |
| TiCCKAR-R | GGGACACATACCAGGACTGC |  |
| TiCCKBR-F | GTCACACTCTGCCTGGTCTC |  |
| TiCCKBR-R | CATAACAGCCGGAGAAGGGG |  |
| TiCCKBRA-F | ACATCCATCAACCCCGAGTG |  |
| TiCCKBRA-R | GAGCAGGATCCGAAGTGTGT |  |

Figure 1- Supplementary Table 1.

Single RNA Guides used for CCKR KO with CRISPR-Cas9 and primers used for real time PCR of tissue distribution in tilapia.

| **Gene** | **Sequence** | **Slope** | **R^2** |
| --- | --- | --- | --- |
| zfEF1A 954F | CTAGCCGTCCCACCGACAAG | -3.2 | 0.99 |
| zfEF1A 1151R | GCAGGCGATGTGAGCAGTGT |  |  |
| zfLHβ 190F | AATGCCTGGTGTTTCAGACC | -3.2 | 1 |
| zfLHβ 353R | AACAGTCGGGCACGTTAATG |  |  |
| zfFSHβ 190F | TGTGGGAGCTGCGTCACAAT | -3.2 | 0.99 |
| zfFSHβ 327R | GCCACGGGGTACACGGAAGAC |  |  |

Figure 5 – Supplementary Table 1.

The primers used for in vivo gene expression analysis in zebrafish (Fig. 4)

Figure 2- supplement movie 1.

Blood flow in the pituitary of live fish. Fish was injected with dextran blue and imaged using two-photon microscopy.

Figure 2- supplement movie 2.

Calcium imaging of LH and FSH cells in vivo, using two-photon microscopy. 2500 frames, 4 Hz image, 150 frames per sec **(A)** Only FSH cells were active; **(B)** both cell types were active. 2000 frames, 10 Hz image, 250 frames per sec. The image to the left is an overlap of the red and green channels showing LH (magenta) and FSH (green) cells.

Figure 3- supplement movie 1.

Calcium imaging of LH and FSH cells *ex vivo*, using confocal microscopy. 2500 frames, 4 Hz image, 150 frames per sec **(A)** LH cell basal activity and GnRH-stimulated calcium wave; **(C)** FSH cell basal activity. **(B)** Another GnRH-stimulated calcium wave of only LH cells; 2500 frames, 4 Hz image, 150 frames per sec. The images to the left is an overlap of the red and green channels showing LH (magenta) and FSH (green) cells.

**Figure4- supplement Movie 1.**

Calcium imaging of LH and FSH cells stimulated with CCK *ex vivo*, using confocal microscopy. 2500 frames, 4 Hz image, 150 frames per sec. The image to the left is an overlap of the red and green channels, showing LH (magenta) and FSH (green) cells.
